# Surveillance of genetic diversity and evolution in locally transmitted SARS-CoV-2 in Pakistan during the first wave of the COVID-19 pandemic

**DOI:** 10.1101/2021.01.13.426548

**Authors:** Muhammad Shakeel, Muhammad Irfan, Zaibunnisa, Muhammad Rashid, Sabeeta Kanwal Ansari, Ishtiaq Ahmad Khan

## Abstract

Surveillance of genetic diversity in the SARS-CoV-2 is extremely important to detect the emergence of more infectious and deadly strains of the virus. In this study, we monitored mutational events in the SARS-CoV-2 genome through whole genome sequencing. The samples (n=48) were collected from the hot spot regions of the metropolitan city Karachi, Pakistan during the four months (May 2020 to August 2020) of first wave of the COVID-19 pandemic. The data analysis highlighted 122 mutations, including 120 single nucleotide variations (SNV), and 2 deletions. Among the 122 mutations, there were 71 singletons, and 51 recurrent mutations. A total of 16 mutations, including 5 nonsynonymous mutations, were detected in spike protein. Notably, the spike protein missense mutation D614G was observed in 31 genomes. The phylogenetic analysis revealed majority of the genomes (36) classified as B lineage, where 2 genomes were from B.6 lineage, 5 genomes from B.1 ancestral lineage and remaining from B.1 sub-lineages. It was noteworthy that three clusters of B.1 sub-lineages were observed, including B.1.36 lineage (10 genomes), B.1.160 lineage (11 genomes), and B.1.255 lineage (5 genomes), which represent independent events of SARS-CoV-2 transmission within the city. The sub-lineage B.1.36 had higher representation from the Asian countries and the UK, B.1.160 correspond to the European countries with highest representation from the UK, Denmark, and lesser representation from India, Saudi Arabia, France and Switzerland, and the third sub-lineage (B.1.255) correspond to the USA. Collectively, our study provides meaningful insight into the evolution of SARS-CoV-2 lineages in spatio-temporal local transmission during the first wave of the pandemic.

## 1.0 Introduction

The novel beta coronavirus, SARS-CoV-2 (Severe acute respiratory syndrome coronavirus 2) which causes respiratory illness named as Covid-19, was first emerged in Wuhan, China in December 2019 and recognized by World Health Organization (WHO) as pandemic in March 2020 (Shereen, Khan, Kazmi, Bashir, and Siddique, 2020). According to John Hopkins University, there were more than 83 million confirmed cases and approximately 1.8 million deaths worldwide by the end of 2020. SARS-CoV-2 is a member of the family Coronaviridae, which comprises many virulent strains that infect humans and animals, including Middle East respiratory syndrome CoV (MERS-CoV) and SARS-CoV (V’kovski, et al., 2020). SARS-CoV-2 is a positive-sense, single-stranded RNA virus with a genome size of approximately 29.8kb. Patients infected with SARS-CoV-2 demonstrate diverse clinical outcomes, ranging from asymptomatic to fatal (Adachi, Koma, Nomaguchi, and Adachi, 2020; Monchatre-Leroy et al., 2017). It contains four major structural proteins including spike (S) protein, membrane (M) protein, and envelope (E) protein, which are embedded in the viral surface envelope, while nucleocapsid (N) protein is in the ribonucleoprotein complex. Furthermore, the viral genome also encodes 16 nonstructural proteins (nsp1-16) and 6 accessory proteins (McBride, Van Zyl, and Fielding, 2014; Hassan, Choudhury, and Roy, 2020). The virus entry in the cell is facilitated by S protein. S1 subunit engages ACE-2 receptor for binding while S protein priming is carried out by binding to cellular serine protease TMPRSS2. This allows the fusion of viral and cellular membranes. This fusion is driven by the S2 subunit of S protein (Hoffmann, et al., 2020). Viruses rapidly evolve during pandemic due to accumulation of mutation. This contributes in viral adaptation, drug resistance and higher transmissibility of more virulent strains (Pachetti et al., 2020). Many descendants of the original Wuhan strain have already been evolved into distinct lineages with potential of vaccine escape. Despite very high mutation rate and rapid emergence of new strains very few mutations have been functionally characterized (Wang, Wang, and Zhuang 2020).

A massive genome sequencing drive is under way globally, to document the genetic diversity of SARS-CoV-2. Several studies has reported large number of mutations in various genes, including S, M, E, N, ORF1ab, ORF3a, ORF6, ORF7, ORF8, and ORF10. It is noteworthy that several regions including nsp1, nsp2 nsp3, nsp12, and nsp15 of ORF1ab, S, as well as ORF8 genes have high mutation rate as compare to other genes (Rahimi, Mirzazadeh, and Tavakolpour, 2020).

Genomic surveillance of a virus after it enters a new population is crucial for designing effective strategies for disease control and prevention (Ladner et al., 2019). The findings of such studies support contact tracing, social distancing, and travel restrictions to contain the spread of SARS-CoV-2. In a study conducted in Northern California from late January to mid-March 2020, using samples from 36 patients spanning nine counties and the Grand Princess cruise ship, phylogenetic analyses revealed the cryptic introduction of at least seven different SARS-CoV-2 lineages into California (Deng *et al*., 2020). Genetic surveillance of COVID-19 studies in Malaysia and other Asian countries highlighted the presence of B.6 lineage in the Asia Pacific region (representing 95% of the world cases of B.6 strains) (Chong *et al*., 2020). The genomic based surveillance of COVID-19 cases in Beijing, China till May 2020 revealed transmission of SARS-CoV-2 in the city via three routes including Wuhan exposure group, foreigner imported cases, and locally transmitted cases (Du *et al*., 2020).

Pakistan, shares borders with world’s most densely populated nations with strong movement of people from and to the hotspots of COVID-19. The country confirmed its first COVID-19 patient on February 26, 2020, in southern city of Karachi. By the end of 2020, there are 479,715 confirmed cases in Pakistan with more than 10,000 deaths. Government adopted progressive disease prevention initiatives to restrict social contacts, reduce the dissemination of viruses and avoid community-based transmissions. Pakistan witnessed peak in June and cases fell from thousands to a few hundred per day in September. Currently, Pakistan is experiencing second wave of infection with around 3000 new cases are diagnosed per day. World Health Organization appreciated overall strategy adopted by Pakistan to successfully contain the virus (WHO 2020 https://www.who.int/news-room/feature-stories/detail/covid-19-in-pakistan-who-fighting-tirelessly-against-the-odds).

The present study attempts to gain insight into the mutational spectrum of SARS-CoV-2 genome in the Pakistani. In this study, we comprehensively analyzed 48 SARS-CoV-2 genome sequences isolated from patients from different hotspots areas of Karachi. Spatiotemporal approach was adopted and samples were collected from May to August 2020. The focus of epidemiological analysis was to identify specific patterns of SARS-CoV-2 transmission through genomic analysis within local population before and during the containment stage of the COVID-19 and compare findings with global data. Finding of this study will help to devise strategies for the future surveillance of potential transmission routes to contain future outbreaks.

## 2.0 Material and Methods

### 2.1 Ethical Consideration, recruitment of patients, and samples collection

The study was approved by the Research Ethics Committee of the International Center for Chemical and Biological Sciences, University of Karachi, and the study design adhered to the ethical considerations according to the Declaration of Helsinki (Helsinki, 2013). For current study, a total of 200 patients were recruited from different hotspot areas of Karachi during the first wave of COVID-19 from May to August 2020. For COVID-19 testing, nasopharyngeal swabs were collected in viral transport medium (VTM) according to the guidelines of the Center for Disease Control and Prevention (CDC, 2020). All the patients were tested positive for COVID-19 by real time PCR using SARS-CoV-2 specific primers and probes at the National Institute of Virology, ICCBS, University of Karachi. Total RNA was isolated from the VTM in a biosafety level-3 (BSL-3) laboratory using QIAamp viral RNA mini kit (Qiagen, Hilden, Germany) following the manufacturer’s protocol. Concentration of the total RNA was evaluated with Qubit fluorometer using Qubit RNA HS assay kit (Thermo Fisher Scientific, MA, USA).

### 2.2 Complementary DNA synthesis

Double stranded complementary DNA (cDNA) was synthesized from the total RNA by using Maxima H Minus Double-Stranded cDNA Synthesis Kit (cat#2561, Thermo Fisher Scientific, MA, USA) according to the manufacturer’s protocol. This involved synthesis of the first strand cDNA followed by the second strand cDNA. For first strand cDNA synthesis, the isolated total RNA and random hexamer primers were incubated at 65°C for 5 minutes followed by addition of 4X First Strand Reaction Mix along with First Strand Enzyme Mix. The reaction mixture was incubated at 25°C for 10 minutes, 50°C for 30 minutes, followed by termination of the reaction at 85°C for 5 minutes. The second strand cDNA synthesis was performed by adding 5X Second Strand Reaction Mix and Second Strand Enzyme Mix to the first strand cDNA synthesis reaction mixture. Final volume was adjusted with nuclease free water, and the reaction mixture was incubated at 16°C for 60 minutes. The reaction was stopped by adding 0.5M EDTA. The double stranded cDNA was purified by using Agencourt AMPure XP beads (Beckman Coulter, CA, USA), and the concentration was evaluated using Qubit DNA HS assay kit (Thermo Fisher Scientific, MA, USA).

### 2.3 DNA library preparation, and whole genome sequencing

A total of 48 samples (male n=29, female n=19) were selected for SARS-CoV-2 whole genome sequencing. Paired-end libraries were constructed from the double stranded cDNA by using Illumina DNA Prep with Enrichment kit (Illumina Inc., San Diego, CA, USA) following the manufacturer’s protocol. Briefly, the tagmentation of purified double stranded cDNA (50 ng) was performed by using bead linked transposomes followed by adapters ligation to the tagmented DNA. Unique indexes (IDT for Illumina DNA/RNA UD Indexes) were added to each tagmented DNA in limited cycles of RCR. The amplified tagmented DNA was purified using sample purification beads. Prior to the enrichment of SARS-CoV-2 genome, the libraries were pooled in 12 plex reactions (multiplexing of 12 samples). DNA fragments of the SARS CoV-2 genome were hybridized with biotinylated respiratory virus oligos (Illumina Inc., San Diego, CA, USA). The DNA fragments hybridized with the custom oligos were captured using streptavidin magnetic beads. The enriched library was amplified followed by purification with Agencourt AMPure XP beads (Beckman Coulter, CA, USA). The concentration of the enriched libraries was determined using Qubit DNA HS assay kit (Thermo Fisher Scientific, MA, USA). The libraries were denatured with 0.2N NaOH followed by dilution to 12 pmole using the hybridization buffer (HT1). Paired-end sequencing (2×75 bases) using MiSeq reaget v2 kit was carried out on Illumina MiSeq (Illumina Inc., San Diego, CA, USA).

### 2.4 Analysis of the sequencing data

The raw data in the binary base call format (.bcl) was converted into fastq format on the MiSeq instrument. Quality of the DNA short reads was assessed using FastQC tool (Andrews, 2010). The short reads were aligned with the reference SARS-CoV-2 genome of the isolate from Wuhan, China (Wuhan-Hu-1 genome, Genbank accession NC_045512) using the BWA-MEM algorithm (Li, 2013). The post alignment processing and variants calling was carried out by using the Samtools package (Dhandapany *et al*., 2009). The samples with coverage ≥88% were processed for downstream analysis. The variants with quality score (QUAL) < 30 were filtered out. The functional annotation of the variants was carried out by using ANNOVAR (Yang *et al*., 2015). For building the whole genome, the consensus sequences were generated from the binary alignment map (bam) files, as described previously (Sah *et al*., 2020). For comparison and validation, DNA short reads were assembled through de novo assembly with Velvet 1.0.0 (Zerbino, 2010) tool using the default parameters. For inferring phylogenetic relationship, the assembled genomes were aligned with the Wuhan-Hu-1 genome using the Muscle multiple sequence alignment tool (Edgar, 2004). The phylogeny was constructed using the RAxML 8.2.12 tool (Stamatakis, 2014) using maximum likelihood algorithm and 100 bootstrap replicates.

## 3.0 Results and Discussion

### 3.1 Description of the Cohort

We selected 48 COVID-19 patients from the public sector hospitals during the peak time of first wave of the pandemic (from 2^nd^ May 2020 to 10^th^ August 2020) in the metropolitan city of Karachi, Pakistan. The samples were included in the study after confirmed diagnosis with real time PCR test from the nasopharyngeal swab specimens. The cohort was comprised of 32 males and 16 females with a median age of 36 years (IQR 23-44 years). The samples were selected from the regions designated as COVID-19 hotspots by the local administrative authority. The disease symptoms were varying in the patients including mild symptoms of low grade fever and flu to moderate symptoms of fever, flue and difficulty in breathing.

### 3.2 Genetic characteristics of the SARS-CoV-2

To uncover the genomic characteristics and diversities of the virus responsible for the COVID-19 in Karachi, deep SARS-CoV-2 genome sequencing was employed using a custom oligos panel designed for the enrichment of respiratory viruses sequences. Through massively parallel sequencing, we obtained 27.112 million paired-end, good quality reads (on average 0.565 million reads per sample). After removing the samples in which viral genome coverage was below 85%, we obtained near complete viral genomes from 37 cases with average depth of coverage of 1371X. After alignment of the short reads with the reference SARS-CoV-2 genome of Wuhan isolate (Wuhan-Hu-1, NC_045512.2), the genetic variations were obtained employing the on-instrument default pipeline of BWA, Samtools, and GATK tools. After filtering out the low quality mutations (those with QUAL<30), there were cumulative 451 mutations at 122 genomic sites (including 120 single nucleotide variation (SNV) sites, and 2 deletion sites) (Supplementary Table S1). These variations include 71 singletons, and 51 recurrent mutations, which appeared more than once within the cohort (Fig.1). Great diversity in mutational sites among the genome assemblies was observed. There were, on average, 12.19 mutational events per genome (SD ±3.57) with a median of 12 mutations per genome (IQR 11-14). The genome-wide nonsynonymous/synonymous ratio was observed as 1.50, which is lower than the precedented nonsyn/syn ratio (1.88) in previous global scale study (van Dorp *et al*., 2020). Further bifurcation indicated that the nonsyn/syn ratio was 1.48 at singleton sites whereas 1.52 at polymorphic sites. Furthermore, the nonsyn/syn ratio at homoplasic sites (those with recurrence in at least 3 genomes) was 1.17. These analyses indicated that the SARS-CoV-2 continued to acquire new nonsynonymous mutational sites within the studied cohort. Next we explored which region was prone to higher number of nonsynonymous mutations. The nonsyn/syn ratio across ORF1ab was 1.75 where the ratio across rest of the genome was 1.23, indicating higher tendency of ORF1ab to acquire mutations at nonsynonymous sites (Fisher Exact P=0.23, OR=1.57, 95% CI=0.671– 3.031).

**Fig 1:**
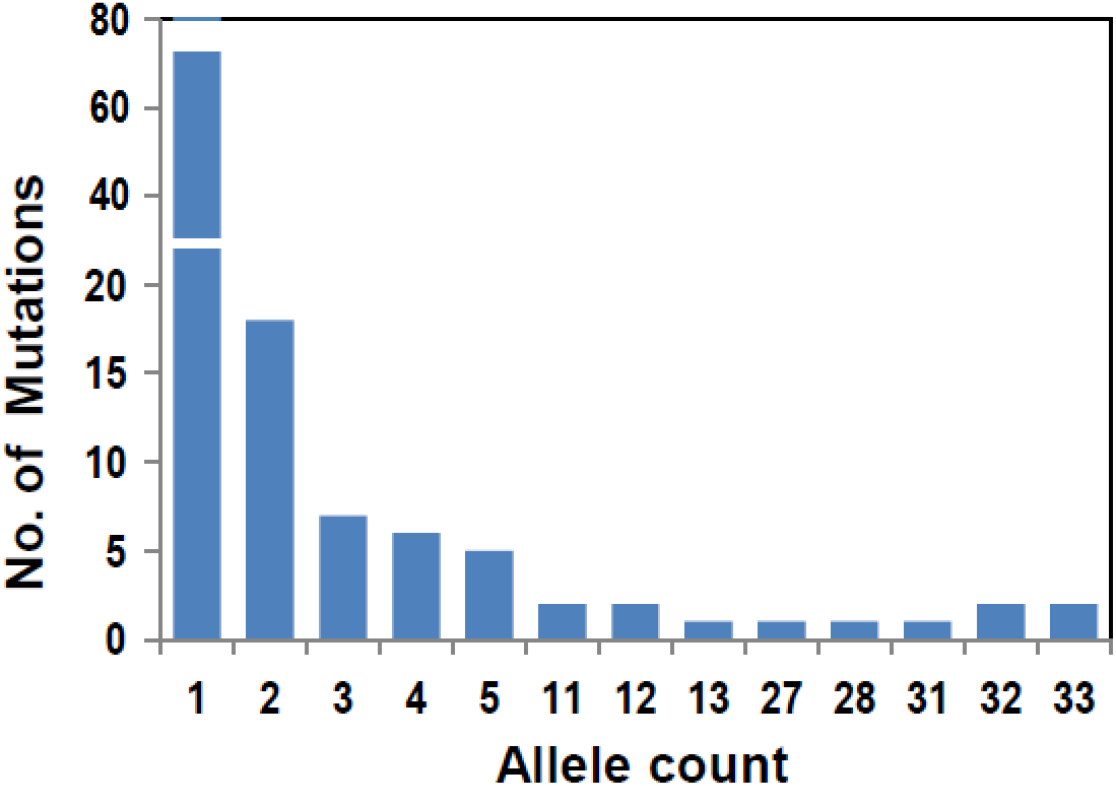
Allelic frequency spectrum of the mutations identified in the SARS-CoV-2 genomes of this study. The larger proportion of mutations comprised of singletons. Notably, highly recurrent mutations (including D614G substitution in the spike protein) were also observed in the analyzed genomes.

### 3.3 Mutational landscape

Meta-analysis of large number of SARS-CoV-2 genomes across multiple countries has demonstrated that one SARS-CoV-2 genome differs from the Wuhan-Wu-I reference strain (NC_045512.2) at maximum of 32 sites (van Dorp *et al*., 2020). We constructed mutational landscape to decipher the genes which are more recurrently mutated than the others. This analysis highlighted that two genes i.e., Spike protein and non-structural protein 3 (nsp3) contained the highest number of recurrent mutations i.e., 35 mutational incidences each (Fig.2). Notably, the spike protein contained highest number of homoplasic sites (6 mutations) including two missense SNVs i.e., the D614G mutation observed in 32 samples (86.5% of the cohort), and Q677H mutation observed in 4 samples (10.8% of the cohort). The other highly recurrent variations included an upstream 5’-UTR mutation 241:C>T (observed in 33 samples), a silent mutation F924F in nsp3 (observed in 31 samples), a missense mutation in nsp12 P4715L (observed in 31 samples), a silent mutation L227L in nsp13 (observed in 12 samples), a silent mutation L280L in nsp14 (observed in 28 samples), silent mutations D294D and G880G in spike protein (observed in 12 and 13 samples respectively), a missense mutation Q57H in orf3a (observed in 31 samples), silent mutation Y71Y and missense mutation H125Y in membrane protein (observed in 27 and 4 samples respectively), a stop-gain mutation E39X in orf7b (observed in 4 samples), and missense mutations S194L and R209I in nucleocapsid protein (observed in 11 samples each). Comparison of the frequency of recurrent mutations in our cohort with the global recurrence showed that the missense mutation V2133A in nsp3 protein was observed in 5 samples of present study cohort, whereas, this mutations has been observed in 2 SARS-CoV-2 genomes in Norway and one in Turkey.

**Fig 2:**
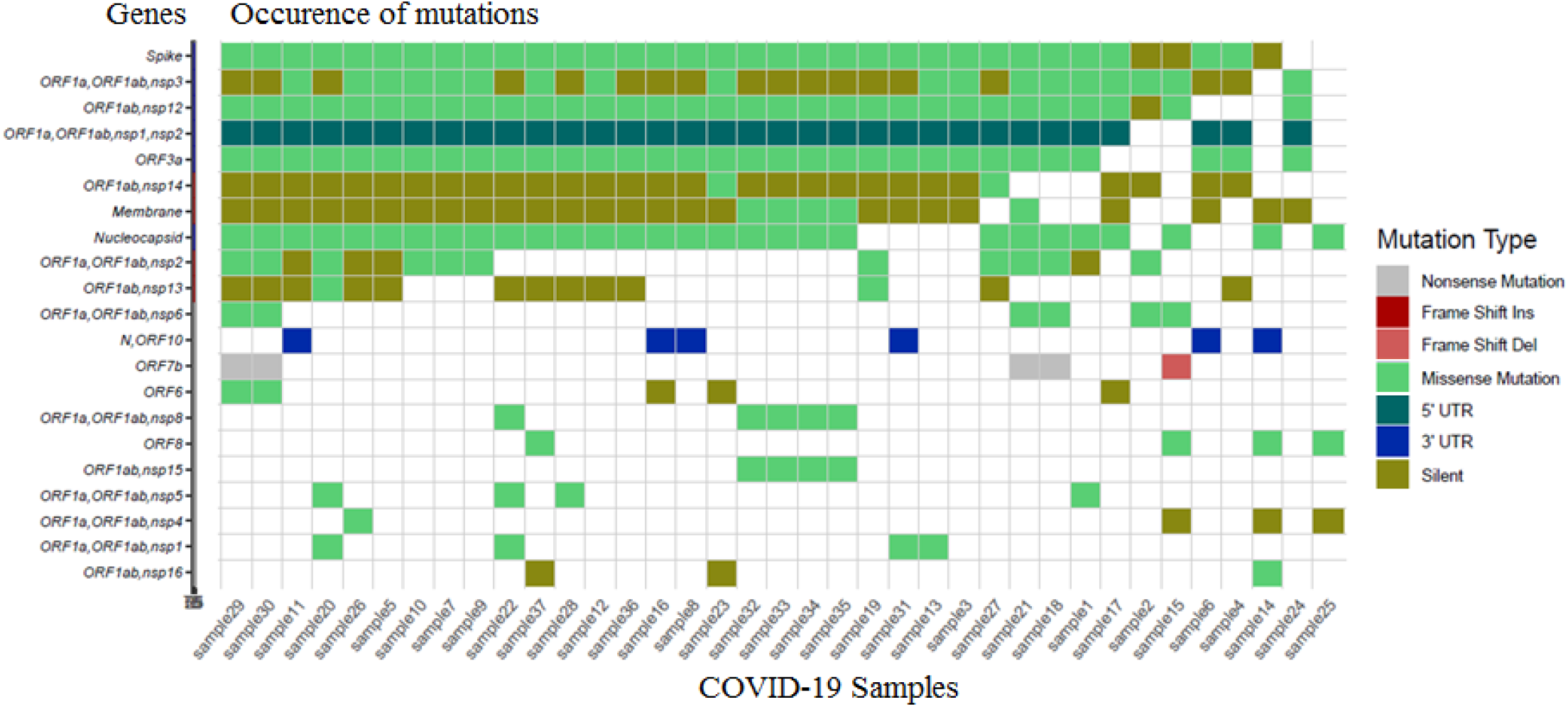
Mutational landscape, recurrence of mutations in genomes of SARS-CoV-2 isolated in Karachi (collected during May 2020 to August 2020)

### 3.4 Spatio-temporal analysis on identified mutations

In order to determine accumulation of mutations in SARS-CoV-2 with the passage of time, we determined mutation rate (total mutations/no. of cases) in each of the four months. This analysis showed that during the month of May 2020, 8.8 mutations per sample were observed, with singletons/recurrent mutations ratio of 2.5. The mutation rate in June 2020 was 10.05 mutations per genome, with singletons/recurrent mutations ratio of 2.65, in July 2020, the mutations per genome were 9.8 with singletons/recurrent mutations ratio of 1.28, and in August 2020, the mutations rate was 12.5 with singletons/recurrent mutations ratio of 0.23 (Fig.3A). Although, there was non-significant difference (one-way ANOVA P > 0.05) in the mutation rate among the four months, yet there was significant increase in recurrent mutations during the month of July and August 2020 (Fig.3B). Next we investigated the mutational rate in genes of ORF1ab, spike, membrane, and nucleocapsid proteins separately (Fig.3C). This analysis indicated that the mutation rate in the ORF1ab during the months of May and June were comparable (5.06 and 5.37 respectively), whereas it decreased in July (4.41) and increased in August (6.37). On the other hand, the mutation rate in the spike protein was 0.99 which increased to almost double in June (1.88), July (1.53), and August (2.09). The mutation rate in the membrane protein was comparable during the four months i.e., 0.78, 0.81, 0.80, 1.30 for May, June, July and August 2020 respectively. Likewise, the mutation rate in the nucleocapsid protein was 1.10, 0.98, 0.91, 1.20 during the months of May, June, July and August 2020 respectively.

**Fig 3:**
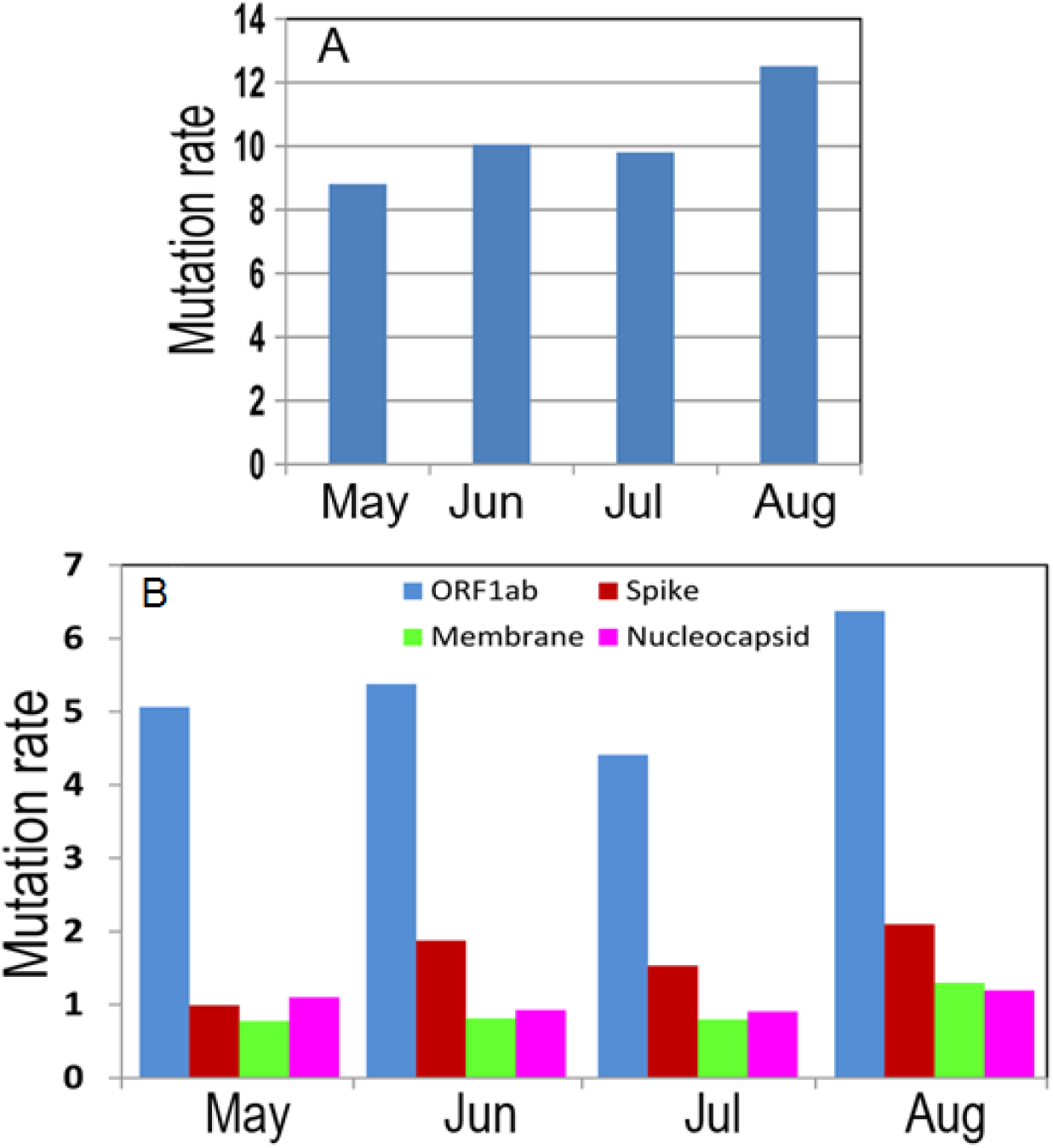
Mutational rate during the four months, May to Aug 2020.

### 3.5 Phylogenetic relationship and lineages in SARS-CoV-2

Given the samples were collected in four different locations of the city, we conducted a phylogenetic analysis to delineate clustering among the SARS-CoV-2 genomes. The phylogenetic analysis revealed that majority of the samples were descendants of B lineages (Fig.4) according to the taxonomic nomenclature proposed by Rambaut *et al*. 2020. Only one genome was found to be of A (Chinese) lineage, whereas, the remaining 36 genomes were of B lineages. Among the genomes of B lineages, one genome was of the parental B lineage and had highest similarity with the Wuhan-Wu-I genome, two genomes were of B.6 (Singapore lineage), whereas remaining 33 were either of B.1 lineage (3 genomes) or further descendants of B.1 lineage. The descendants of B.1 comprised of three types of high prevalent lineages i.e., B.1.36 (10 genomes), B.1.160 (11 genomes), and B.1.255 (5 genomes). These three types of B.1 sub-lineages seem to be spread in the city during independent transmission events. The B.1.36 is global lineage with lots of representation of sequences from the India, Saudi Arabia, and UK (Ishtiaque *et al*., 2020; Joshi *et al*., 2020). The B.1.160 lineage is the most recently split from B.1.36 lineage and has mostly been reported from the European countries with major representation in the UK followed by Denmark, France and Switzerland (Rambaut *et al*., 2020). The third major lineage in our genomes B.1.255 has major global representation with significantly higher representation in the USA. Notably, one genome was observed from B.1.184 lineage, which has 100% representation in India (Andrew *et al*., 2020). These findings were validated in the hierarchical clustering where three clusters of genomes were observed (Fig.5), where the founding lineages (B, B.1) appeared in the center of the principle component analysis (PCA) plot, whereas, the three sub-lineages clustered at the peripheries of the plot.

**Fig 4:**
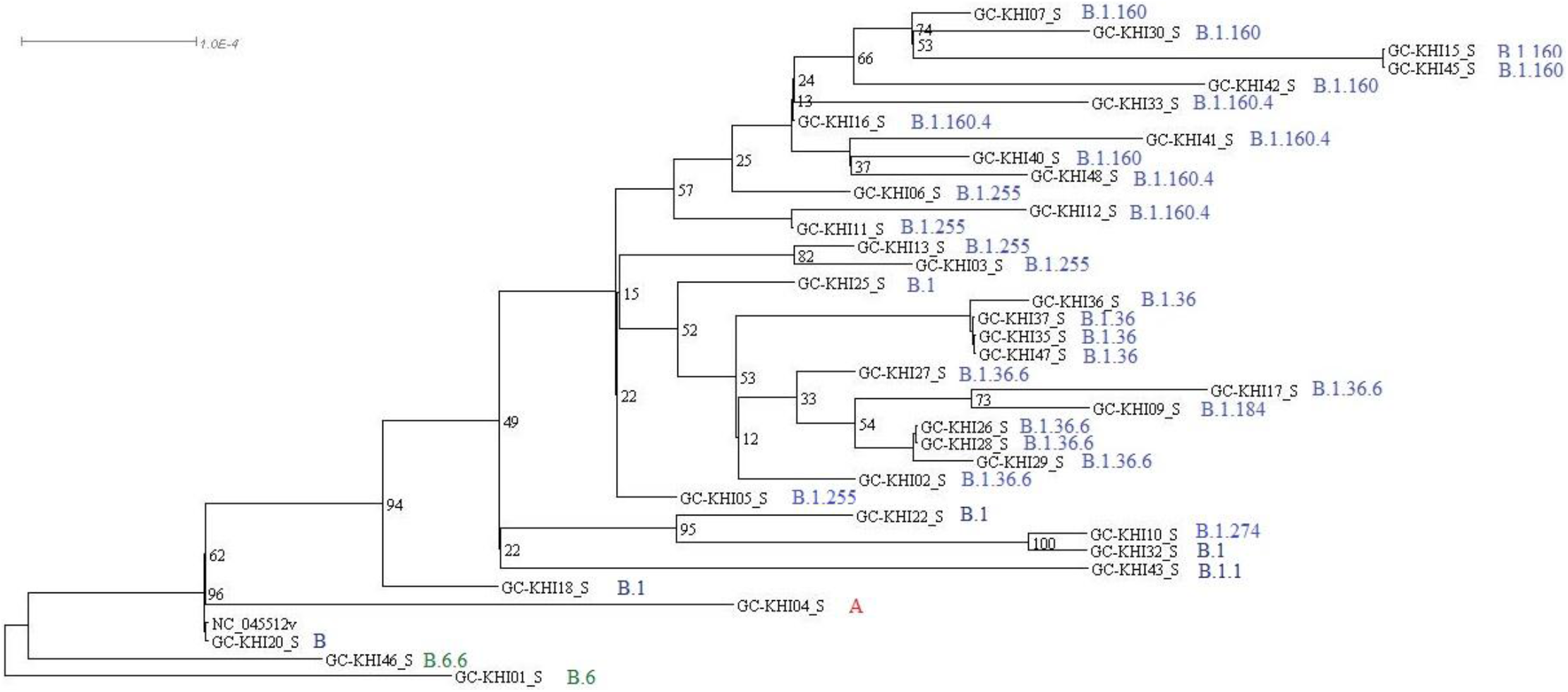
Phylogenetic analysis of the SARS-CoV-2 genomes. The analysis indicated three types of lineages B.1.36, B.1.160, and B.1.255 in the analyzed cohort.

**Fig 5:**
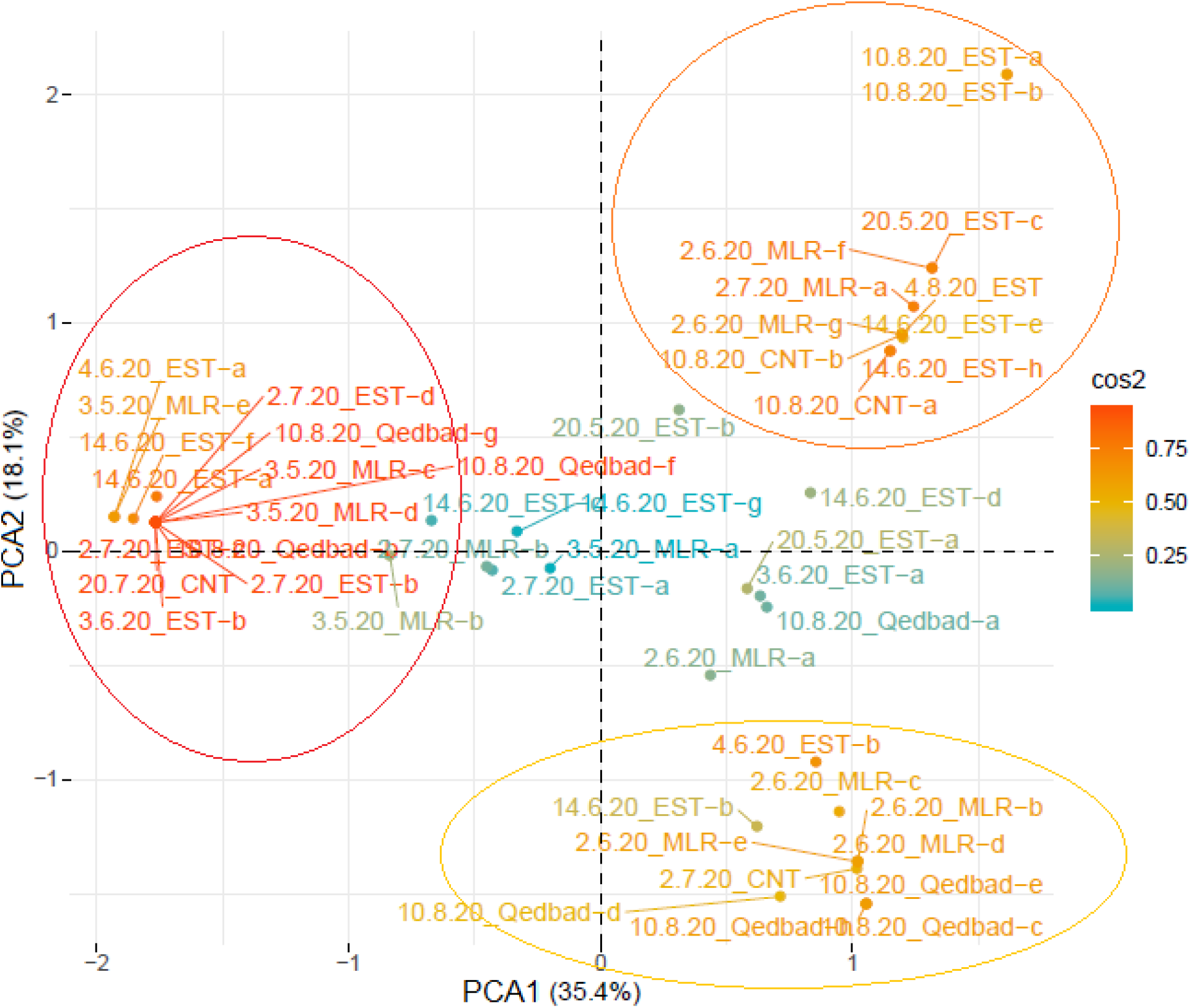
Hierarchal clustering of the SARS-CoV-2 genomes of this study using the mutational events in genomes. The genomes of B.1 parental lineages appeared in the center of the PCA plot, whereas, the three sub-lineages clustered at the peripheries.

### Conclusion

To the best of our knowledge this study presents first comprehensive report on the surveillance of genomic evolution in the SARS-CoV-2 transmitted locally in the metropolitan city of Karachi, Pakistan during the four months of first wave of the COVID-19 pandemic. The schematic analysis provided meaningful insight into the lineages being transmitted within the city. The higher prevalence of three types of B.1 sub-lineages highlighted that the virus most likely had entered into the region through the travellers from the Europe, USA, Saudi Arabia, and India. The findings of this study enabled to monitor whether a more deadly strain of SARS-CoV-2 is being spreading in the country or else. These preliminary analyses present mutational and lineages status of the genomes, further analyses are underway to find the impact of identified mutations on viral infectivity and fatality.

### Data Submission

The whole genome sequence data reported in this paper have been deposited in the Genome Warehouse in National Genomics Data Center (Genome Warehouse, 2021), Beijing Institute of Genomics (China National Center for Bioinformation), Chinese Academy of Sciences, under BioProject accession number PRJCA004109 (publicly accessible at https://bigd.big.ac.cn/gwh), and NCBI Genbank accession numbers MW447609-MW447645.

## Acknowledgement

This study was conducted through the funding provided by the host institution, Dr. Panjwani Center for Molecular Medicine and Drug Research (PCMD), ICCBS, University of Karachi, Pakistan.

---

**Supplementary Table S1:** The recurrent and singleton mutations identified in the SARS-CoV-2 genomes of present study

| S.No. | Mutation | Gene | Mutation class | Mutation occurrence |
| --- | --- | --- | --- | --- |
| 1 | C241T | ORF1a,ORF1ab,nsp1,nsp2 | upstream | 33 |
| 2 | P.Q63L | ORF1a,ORF1ab,nsp1 | nonsynonymous_SNV | 1 |
| 3 | P.G82S | ORF1a,ORF1ab,nsp1 | nonsynonymous_SNV | 2 |
| 4 | P.V116L | ORF1a,ORF1ab,nsp1 | nonsynonymous_SNV | 1 |
| 5 | P.N188N | ORF1a,ORF1ab,nsp2 | synonymous_SNV | 1 |
| 6 | P.K261N | ORF1a,ORF1ab,nsp2 | nonsynonymous_SNV | 2 |
| 7 | P.F275L | ORF1a,ORF1ab,nsp2 | nonsynonymous_SNV | 3 |
| 8 | P.R287G | ORF1a,ORF1ab,nsp2 | nonsynonymous_SNV | 2 |
| 9 | P.Q362Q | ORF1a,ORF1ab,nsp2 | synonymous_SNV | 4 |
| 10 | P.H374H | ORF1a,ORF1ab,nsp2 | synonymous_SNV | 1 |
| 11 | P.S481F | ORF1a,ORF1ab,nsp2 | nonsynonymous_SNV | 1 |
| 12 | P.T592I | ORF1a,ORF1ab,nsp2 | nonsynonymous_SNV | 1 |
| 13 | P.V627F | ORF1a,ORF1ab,nsp2 | nonsynonymous_SNV | 2 |
| 14 | P.Y717Y | ORF1a,ORF1ab,nsp2 | synonymous_SNV | 3 |
| 15 | P.F924F | ORF1a,ORF1ab,nsp3 | synonymous_SNV | 32 |
| 16 | P.G934C | ORF1a,ORF1ab,nsp3 | nonsynonymous_SNV | 1 |
| 17 | P.T999I | ORF1a,ORF1ab,nsp3 | nonsynonymous_SNV | 1 |
| 18 | P.L1175I | ORF1a,ORF1ab,nsp3 | nonsynonymous_SNV | 1 |
| 19 | P.Y1354Y | ORF1a,ORF1ab,nsp3 | synonymous_SNV | 1 |
| 20 | P.K1396R | ORF1a,ORF1ab,nsp3 | nonsynonymous_SNV | 1 |
| 21 | P.A1485A | ORF1a,ORF1ab,nsp3 | synonymous_SNV | 1 |
| 22 | P.T1754I | ORF1a,ORF1ab,nsp3 | nonsynonymous_SNV | 1 |
| 23 | P.F1925F | ORF1a,ORF1ab,nsp3 | synonymous_SNV | 2 |
| 24 | P.D1926G | ORF1a,ORF1ab,nsp3 | nonsynonymous_SNV | 1 |
| 25 | P.S2015R | ORF1a,ORF1ab,nsp3 | nonsynonymous_SNV | 1 |
| 26 | P.T2016K | ORF1a,ORF1ab,nsp3 | nonsynonymous_SNV | 2 |
| 27 | P.V2133A | ORF1a,ORF1ab,nsp3 | nonsynonymous_SNV | 5 |
| 28 | P.T2408I | ORF1a,ORF1ab,nsp3 | nonsynonymous_SNV | 3 |
| 29 | P.K2497E | ORF1a,ORF1ab,nsp3 | nonsynonymous_SNV | 1 |
| 30 | P.L2564F | ORF1a,ORF1ab,nsp3 | nonsynonymous_SNV | 1 |
| 31 | P.A2618V | ORF1a,ORF1ab,nsp3 | nonsynonymous_SNV | 1 |
| 32 | P.Q2702H | ORF1a,ORF1ab,nsp3 | nonsynonymous_SNV | 3 |
| 33 | P.S2839S | ORF1a,ORF1ab,nsp4 | synonymous_SNV | 1 |
| 34 | P.T2846I | ORF1a,ORF1ab,nsp4 | nonsynonymous_SNV | 1 |
| 35 | P.Y3098Y | ORF1a,ORF1ab,nsp4 | synonymous_SNV | 1 |
| 36 | P.L3293F | ORF1a,ORF1ab,nsp5 | nonsynonymous_SNV | 1 |
| 37 | P.K3353R | ORF1a,ORF1ab,nsp5 | nonsynonymous_SNV | 1 |
| 38 | P.P3447S | ORF1a,ORF1ab,nsp5 | nonsynonymous_SNV | 1 |
| 39 | P.V3475F | ORF1a,ORF1ab,nsp5 | nonsynonymous_SNV | 1 |
| 40 | P.L3606F | ORF1a,ORF1ab,nsp6 | nonsynonymous_SNV | 4 |
| 41 | P.T3646A | ORF1a,ORF1ab,nsp6 | nonsynonymous_SNV | 2 |
| 42 | P.A3657T | ORF1a,ORF1ab,nsp6 | nonsynonymous_SNV | 2 |
| 43 | P.A3956V | ORF1a,ORF1ab,nsp8 | nonsynonymous_SNV | 4 |
| 44 | P.P4075S | ORF1a,ORF1ab,nsp8 | nonsynonymous_SNV | 1 |
| 45 | P.N4235N | ORF1a,ORF1ab,nsp9 | synonymous_SNV | 1 |
| 46 | P.A4489V | ORF1ab,nsp12 | nonsynonymous_SNV | 1 |
| 47 | P.C4698C | ORF1ab,nsp12 | synonymous_SNV | 1 |
| 48 | P.P4715L | ORF1ab,nsp12 | nonsynonymous_SNV | 32 |
| 49 | P.F4807F | ORF1ab,nsp12 | synonymous_SNV | 2 |
| 50 | P.A4921V | ORF1ab,nsp12 | nonsynonymous_SNV | 2 |
| 51 | P.V4979L | ORF1ab,nsp12 | nonsynonymous_SNV | 1 |
| 52 | P.H4991H | ORF1ab,nsp12 | synonymous_SNV | 1 |
| 53 | P.L5028L | ORF1ab,nsp12 | synonymous_SNV | 2 |
| 54 | P.L5167L | ORF1ab,nsp12 | synonymous_SNV | 1 |
| 55 | P.T127I | ORF1ab,nsp13 | nonsynonymous_SNV | 2 |
| 56 | P.L227L | ORF1ab,nsp13 | synonymous_SNV | 12 |
| 57 | P.L256L | ORF1ab,nsp13 | synonymous_SNV | 1 |
| 58 | P.V495V | ORF1ab,nsp13 | synonymous_SNV | 1 |
| 59 | P.M49I | ORF1ab,nsp14 | nonsynonymous_SNV | 1 |
| 60 | P.L157F | ORF1ab,nsp14 | nonsynonymous_SNV | 1 |
| 61 | P.L280L | ORF1ab,nsp14 | synonymous_SNV | 28 |
| 62 | P.S454S | ORF1ab,nsp14 | synonymous_SNV | 1 |
| 63 | P.L495L | ORF1ab,nsp14 | synonymous_SNV | 1 |
| 64 | P.V148F | ORF1ab,nsp15 | nonsynonymous_SNV | 4 |
| 65 | P.V104V | ORF1ab,nsp16 | synonymous_SNV | 1 |
| 66 | P.R216R | ORF1ab,nsp16 | synonymous_SNV | 1 |
| 67 | P.M260I | ORF1ab,nsp16 | nonsynonymous_SNV | 1 |
| 68 | P.D80D | Spike | synonymous_SNV | 1 |
| 69 | P.D294D | Spike | synonymous_SNV | 12 |
| 70 | P.T302T | Spike | synonymous_SNV | 1 |
| 71 | P.S305S | Spike | synonymous_SNV | 5 |
| 72 | **P.D614G** | Spike | nonsynonymous_SNV | 33 |
| 73 | P.D663D | Spike | synonymous_SNV | 1 |
| 74 | P.Q677H | Spike | nonsynonymous_SNV | 4 |
| 75 | P.Y789Y | Spike | synonymous_SNV | 2 |
| 76 | P.G880G | Spike | synonymous_SNV | 13 |
| 77 | P.T887T | Spike | synonymous_SNV | 1 |
| 78 | P.I1018I | Spike | synonymous_SNV | 2 |
| 79 | P.F1148F | Spike | synonymous_SNV | 5 |
| 80 | P.D1153G | Spike | nonsynonymous_SNV | 1 |
| 81 | P.P1162S | Spike | nonsynonymous_SNV | 2 |
| 82 | P.D1260D | Spike | synonymous_SNV | 1 |
| 83 | P.V1264L | Spike | nonsynonymous_SNV | 3 |
| 84 | P.I20V | ORF3a | nonsynonymous_SNV | 1 |
| 85 | P.K21N | ORF3a | nonsynonymous_SNV | 1 |
| 86 | P.Q57H | ORF3a | nonsynonymous_SNV | 31 |
| 87 | P.H78H | ORF3a | synonymous_SNV | 1 |
| 88 | P.T223I | ORF3a | nonsynonymous_SNV | 1 |
| 89 | P.S4S | Membrane | synonymous_SNV | 5 |
| 90 | P.L56L | Membrane | synonymous_SNV | 1 |
| 91 | P.Y71Y | Membrane | synonymous_SNV | 27 |
| 92 | P.H125Y | Membrane | nonsynonymous_SNV | 4 |
| 93 | P.T130T | Membrane | synonymous_SNV | 1 |
| 94 | P.T175M | Membrane | nonsynonymous_SNV | 1 |
| 95 | P.I32I | ORF6 | synonymous_SNV | 1 |
| 96 | P.L44V | ORF6 | nonsynonymous_SNV | 2 |
| 97 | P.D61D | ORF6 | synonymous_SNV | 2 |
| 98 | P.F13fs | ORF7b | frameshift_deletion | 1 |
| 99 | P.L25F | ORF7b | nonsynonymous_SNV | 1 |
| 100 | P.E39X | ORF7b | stopgain | 5 |
| 101 | P.W45L | ORF8 | nonsynonymous_SNV | 1 |
| 102 | P.L84S | ORF8 | nonsynonymous_SNV | 1 |
| 103 | P.Q91L | ORF8 | nonsynonymous_SNV | 1 |
| 104 | P.P13L | Nucleocapsid | nonsynonymous_SNV | 1 |
| 105 | P.R40L | Nucleocapsid | nonsynonymous_SNV | 2 |
| 106 | P.Y111Y | Nucleocapsid | synonymous_SNV | 1 |
| 107 | P.S194L | Nucleocapsid | nonsynonymous_SNV | 11 |
| 108 | P.S202N | Nucleocapsid | nonsynonymous_SNV | 1 |
| 109 | P.R203K | Nucleocapsid | nonsynonymous_SNV | 1 |
| 110 | P.R203R | Nucleocapsid | synonymous_SNV | 1 |
| 111 | P.G204R | Nucleocapsid | nonsynonymous_SNV | 1 |
| 112 | P.R209I | Nucleocapsid | nonsynonymous_SNV | 11 |
| 113 | P.F314F | Nucleocapsid | synonymous_SNV | 3 |
| 114 | P.S327L | Nucleocapsid | nonsynonymous_SNV | 1 |
| 115 | P.K375K | Nucleocapsid | synonymous_SNV | 1 |
| 116 | P.T379I | Nucleocapsid | nonsynonymous_SNV | 1 |
| 117 | P.D402Y | Nucleocapsid | nonsynonymous_SNV | 1 |
| 118 | P.Y14C | ORF10 | nonsynonymous_SNV | 1 |
| 119 | G29742A | N,ORF10 | downstream | 1 |
| 120 | GAGTGTAC29755G | N,ORF10 | downstream | 1 |
| 121 | C29838T | N,ORF10 | downstream | 1 |
| 122 | C29870A | N,ORF10 | downstream | 3 |

